# Nucleotide Excision Repair of Aflatoxin-induced DNA Damage within the 3D Human Genome Organization

**DOI:** 10.1101/2023.09.27.559858

**Authors:** Yiran Wu, Muhammad Muzammal Adeel, Aziz Sancar, Wentao Li

## Abstract

Aflatoxin B1 (AFB1), a potent mycotoxin, is one of the two primary risk factors that cause liver cancer. In the liver, the bioactivated AFB1 intercalates into the DNA double helix to form a bulky DNA adduct which will lead to mutation if left unrepaired. We have adapted the tXR-seq method to measure the nucleotide excision repair of AFB1-induced DNA adducts. We have found that transcription-coupled repair plays a major role in the damage removal process and the released excision products have a distinctive length distribution pattern. We further analyzed the impact of 3D genome organization on the repair of AFB1-induced DNA adducts. We have revealed that chromosomes close to the nuclear center and A compartments undergo expedited repair processes. Notably, we observed an accelerated repair around both TAD boundaries and loop anchors. These findings provide insights into the complex interplay between repair, transcription, and 3D genome organization, shedding light on the mechanisms underlying AFB1-induced liver cancer.

**TEASER:** Genome-wide mapping repair of AFB1-caused DNA damage by tXR-seq reveals 3D genome organization’s significant impact on the repair.

## INTRODUCTION

Aflatoxins, discovered in the early 1960’s (*1*), are a group of mycotoxins produced by certain fungal species such as *Aspergillus flavus* and *Aspergillus parasiticus*. These fungi can contaminate crops and food, especially in regions with warm and humid climates. Aflatoxin B1 (AFB1) is the most prevalent and highly carcinogenic member of this mycotoxin family. Chronic dietary exposure to aflatoxin and hepatitis B virus infection are the two primary risk factors associated with hepatocellular carcinoma (HCC), which is the third leading cause of global cancer-related fatalities (*2*). Aflatoxin initiates hepatocarcinogenesis through genotoxic processes, including metabolic activation to an epoxide, formation of the aflatoxin-DNA adducts, and mutagenesis. Upon ingestion, AFB1 is absorbed and transported to the liver, where it is metabolized into AFB1-8,9-epoxide by cytochrome-P450 enzymes. This reactive epoxide intercalates into the DNA double helix and covalently bonds with the most nucleophilic N^7^ atom of deoxyguanosine to form an AFB1-N^7^-dG bulky adduct. This resulting AFB1-N^7^-dG adduct is unstable and can undergo depurination or spontaneous ring-opening hydrolysis which leads to the formation of an AFB1-formamidopyrimidine-dG (AFB1-FAPY-dG) adduct. Compared to AFB1-N^7^-dG, the ring-opened AFB1-FAPY-dG is more flexible and stable (*3*). Duplex DNA containing AFB1-FAPY-dG is less distorted and has a lower repair propensity than the AFB1-N^7^-dG adducted DNA, which might contribute to the higher mutagenicity of AFB1-FAPY-dG DNA adduct (*4*). If left unrepaired, the two types of AFB1-dG adduct can be bypassed during replication by error-prone translesion synthesis (TLS) DNA polymerases, leading to predominantly G to T transversions (*5, 6*). A particularly striking example is the mutation hotspot at codon 249 (AGG to AGT) of the *TP53* gene, commonly observed in HCC patients from regions with high risk of dietary AFB1 exposure (*7*). Computational analysis of somatic mutations from HCC patients with AFB1 exposure reveals a unique mutation pattern known as COSMIC mutational signature 24, which has a strong transcriptional strand bias (*8, 9*). Besides targeting duplex DNA, AFB1-8,9 epoxide can be hydrolyzed to AFB1 dialdehyde, which reacts with lysine’s amino group in serum albumin to form protein adducts (*10*). The AFB1-albumin adduct in human blood samples is a facile biomarker for assessing chronic dietary AFB1 exposure (*11*). Previous studies suggest that reactive oxygen species generated during AFB1 metabolism in liver cells initiate lipid peroxidation and its byproducts, acetaldehyde and crotonaldehyde, can also damage DNA by forming bulky cyclic propano-dG adducts (*12, 13*). The sequence specificity of this type of damage formation and repair may shape the mutation hotspot observed at codon 249 of the *TP53* gene.

Nucleotide excision repair is a highly conserved and versatile repair pathway that removes various types of helix distorting and bulky DNA lesions, including AFB1-dG adducts, UV-induced cyclobutane pyrimidine dimers (CPDs) and pyrimidine-pyrimidone (6–4) photoproducts [(6–4)PPs], and benzo[a]pyrene (BaP)-induced DNA adducts (*14*). NER comprises two subpathways: global genomic repair and transcription-coupled repair (TCR). While global genomic repair removes DNA damage throughout the entire genome, TCR, triggered by the stalling of elongating RNA polymerase II (RNAPII) at a DNA lesion (*15*), is dedicated to the faster repair of DNA lesions in the transcribed strand (TS) than in the non-transcribed strand (NTS) (*16, 17*). The two subpathways differ only at the damage recognition step and share the same DNA repair machinery in the remaining repair processes including dual incisions bracketing the lesion, release of the excision products, gap filling, and ligation. Interestingly, even though the dual-incision mechanism is quite similar for prokaryotes, archaea, and eukaryotes, the pattern of dual-incision varies among different species. In *E. coli* and *M. smegmatis*, dual incisions occur 7 nucleotide (nt) 5’ and 3-4 nt 3’ to the UV damage site to generate 12-13 nt long excision products (*18*). In the archaea, *M. thermoautotrophicum*, 5’ incision is only 5-6 nt to the UV damage resulting in the 11-nt-long excision product (*19*). In yeast cells, the dual incision sites are at 13-18 nt 5′ and 6-7 nt 3′ to the damage and the predominant excision products are 23-24 nt in length (*20*). In humans, Drosophila melanogaster, and Arabidopsis thaliana, the UV damage is excised by dual incisions 19-22 nt 5′ and 5-6 nt 3′ to the damage site yielding predominantly 26-27-nt excision products (*21–23*). Nucleotide excision repair is generally presumed to be the sole repair pathway that can remove the AFB1-dG adducts (*24, 25*), whereas recent studies show that the DNA glycosylase NEIL1 recognizes and excises the AFB1-FAPY-dG in a sequence-dependent manner, suggesting base excision repair plays a role in the removal of AFB1-FAPY-dG adducts as well (*26, 27*).

Three-dimensional (3D) genome organization plays a crucial role in various nuclear processes, including gene regulation, DNA replication, and DNA damage and repair. Within the cell nucleus, the entire genome is organized into distinct hierarchical structures at various length scales, including chromosome territories, A/B compartments, topologically associating domains (TADs), and chromatin loops. Each chromosome occupies a distinct territory within the nucleus, which is termed a chromosome territory. At the megabase scale, chromosomes are divided into A and B compartments which represent active and inactive chromatin regions, respectively. TAD refers to a specific self-interacting region, in which chromatin regions interact more frequently than the neighboring regions (*28*). Chromatin interactions, particularly those involving enhancer-promoter interactions during gene regulation, are facilitated by a process known as loop extrusion. This mechanism allows distant regulatory elements, such as promoters and enhancers, to come into spatial proximity of targeted genes. In theory, higher-order chromatin organizations can affect DNA damage distribution and repair efficiency. Meanwhile, DNA damage formation, DNA damage response, and repair events can alter the 3D genome structure (*29, 30*). However, how the multiple levels of genomic organizations interplay with DNA damage formation and repair remain largely unexplored.

With the advent of high throughput next-generation sequencing (NGS) techniques, a variety of NGS-based methods, such as high-throughput chromosome conformation capture (Hi-C), Chromatin Interaction Analysis with Paired-End-Tag sequencing (ChIA-PET), Damage-sequencing (Damage-seq), and eXcision Repair-sequencing (XR-seq), have been invented to investigate 3D genome structures, DNA damage formation, and DNA repair (*31, 32*). To date, genome-wide nucleotide excision repair of different DNA damaging agents, such as UV, BaP, cisplatin, oxaliplatin, and 5-ethynyl-2′-deoxyuridine (EdU), have been analyzed at single nucleotide resolution by using XR-seq in a variety of organisms (*21, 33–35*). However, how DNA damage caused by AFB1, the most potent hepatocarcinogen, is removed by nucleotide excision repair at single-base resolution on a genome-wide scale is still unexplored. Furthermore, how the hierarchical levels of 3D genome organization affect the repair of AFB1-dG adducts remains unknown.

In this study, we adapted the translesion XR-seq (tXR-seq) method (*33*) to map AFB1-dG repair in the entire human genome at single-base resolution. Our results showed that AFB1-dG adducts are mainly removed by TCR and the excision products have a unique length distribution pattern. To gain a comprehensive understanding of the AFB1-dG repair, we then integrated datasets from Hi-C, ChIA-PET, DNA fluorescence *in situ* hybridization (FISH), and tXR-seq to investigate the AFB1-dG repair across four different organization levels, including chromosome territories, A/B compartments, TADs, and chromatin loops. Our analysis revealed a heterogeneous AFB1-dG repair landscape within the 3D genome organization. These findings provide insights into the complex interplay between nucleotide excision repair, transcription, and 3D genome organization, paving the way for our understanding of AFB1-induced HCC.

## RESULTS

### Adaptation of tXR-seq for genome-wide mapping nucleotide excision repair of AFB1- induced DNA damage

During nucleotide excision repair, excision products containing the damage are released in complex with TFIIH-XPG. In the conventional XR-seq method (*21*), these excision products are captured, sequenced, and mapped to the reference genome. Before library amplification by PCR, the DNA damage within the excision products must be removed either enzymatically or chemically. In contrast, tXR-seq employs appropriate translesion DNA synthesis (TLS) polymerases to bypass the damage before PCR amplification, making it applicable for essentially all types of DNA lesions removed by nucleotide excision repair (*33*). As the quantity of excision products is crucial for the success of tXR-seq, we performed an *in vivo* excision assay using HepG2 cells, a commonly studied cell line, before the starting of AFB1-dG tXR-seq procedure (Fig. 1A). As shown in Fig. 1B, although the yield is lower than that in UVC-treated (20 J/m^2^) HepG2 cells, the excision products are detectable when HepG2 cells are treated with AFB1 (40 µM) for 4 hrs.

**Fig. 1.**
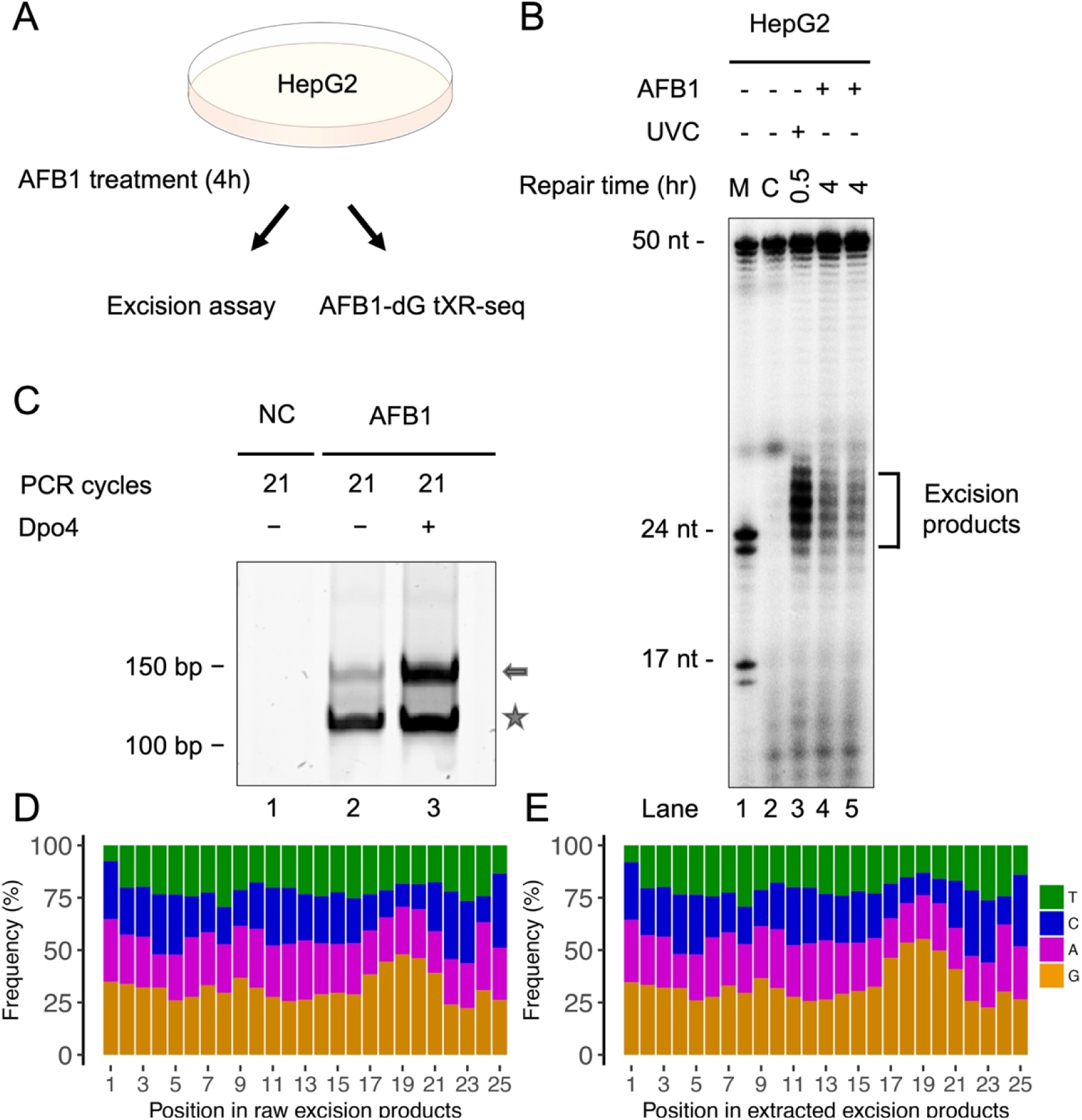
*In vivo* excision assay and analyses of single-nucleotide frequencies for the excision products obtained from AFB1-dG tXR-seq. (A) Overview of the AFB1 *in vivo* excision assay and tXR-seq procedure. (B) Detection of excision products by excision assay after UVC (20 J/m^2^) or AFB1 (40 µM) treatment in HepG2 cells. Excision products from one-quarter of a 150 mm petri dish of UVC-treated HepG2 cells were loaded into the gel, while for AFB1-treated HepG2 cells, excision products from two 150 mm petri dishes were used in the excision assay. M, DNA size marker; C, control. (C) Analysis of the dsDNA library for AFB1-dG tXR-seq through 10% native polyacrylamide gel electrophoresis, with the right arrow indicating PCR products with inserts and the asterisk indicating adaptor-only PCR products. NC, non-template control. (D) Single-nucleotide frequencies for the 25-mers in the raw reads from AFB1-dG tXR-seq. (E) Single-nucleotide frequencies for the 25-mers in the reads extracted from the human reference genome.

For the AFB1-dG tXR-seq, we converted AFB1-N^7^-dG adducts to ring-opened AFB1-FAPY-dG adducts after capturing the excision products by TFIIH/XPG immunoprecipitation (IP). The unstable AFB1-N^7^-dG adducts are quantitatively abundant within 24 hrs after AFB1 treatment (*36*) and the conversion avoids their depurination, which would prevent their detection in the following tXR-seq procedures. We then determined the optimal AFB1 antibody and TLS polymerase that could capture the excision products and bypass the converted AFB1-FAPY-dG lesion, respectively (fig. S1). Gel electrophoresis of the libraries, generated by PCR amplification of the AFB1-FAPY-dG containing excision products, indicated that sulfolobus DNA polymerase IV (Dpo4) could bypass the AFB1-FAPY-dG adduct and be used for our tXR-seq method (Fig. 1C and fig. S1).

After sequencing, the nucleotide frequencies of the excision products from the raw reads were analyzed. As shown in Figure 1 D, guanines (Gs) are enriched at positions 17-22 for 25-mer excision products. Since the Y-family DNA polymerase Dpo4 introduces mainly G to T transversions during lesion bypass, the raw sequencing reads from our AFB1-dG tXR-seq may contain these mutations. After mapping the raw reads to the human reference genome, we then extracted the mapped reads sequences from the reference genome and compared them with our raw sequenced reads. As expected, Gs frequencies at positions 17-22 for the 25-mer excision products in the extracted reads shown in Fig. 1E are higher than in the raw reads (Fig. 1D). Further analysis of the frequency change after bypass confirmed the predominant G to T transversions introduced by the error-prone Dpo4 (fig. S2).

### Unique length distribution of excision products from AFB1-dG repair

Excision products released during nucleotide excision repair vary in size among different species, with *E. coli*, yeast, and humans producing predominant excision products of 12-13 nt, 23-24 nt, and 26-27 nt, respectively (*18, 20*). Bulky DNA adducts induced by different DNA damaging agents can distort the DNA double helix to various degrees, affecting their recognition by the excision repair machinery (*37*). AFB1-N^7^-dG and AFB1-FAPY-dG adducts form intercalated conformations in the DNA double helix, contributing to their repair resistance (*38*). However, it remains unknown whether these two types of DNA lesions affect the length distribution of excision products.

In our analysis of AFB1-dG repair in HepG2 cells, we observed a distinct excision product length distribution, with the predominant length being only 25 nt, in contrast to excision products containing other DNA damaging agents-induced adducts where 26-mers are most frequent (Fig. 2, B to F) (*21, 33–35*). Typically, after dual incision, excision products are released in a stable complex with TFIIH, protecting them from degradation by cellular nucleases such as TREX1 (*39, 40*). The TFIIH immunoprecipitation step in the XR-seq and tXR-seq protocols captures excision products with maximum length, excluding those released from TFIIH and partially degraded. The shorter predominant excision product length in AFB1-dG repair may be attributed to the relatively large size of the AFB1-dG bulky adduct, as supported by the length distribution of excision products from repair of the large-sized BPDE-dG adduct (Fig. 2B). Our nucleotide frequency analysis across 24-mers to 27-mers from AFB1-dG repair consistently shows a peak in G frequency at position 19, suggesting dual incision occurs 18-20 nt 5’ and 5-6 nt 3’ to the lesion (fig. S3). These findings offer insights into the unique dynamics of AFB1- induced DNA lesion repair and its impact on the excision products’ length distribution.

**Fig. 2.**
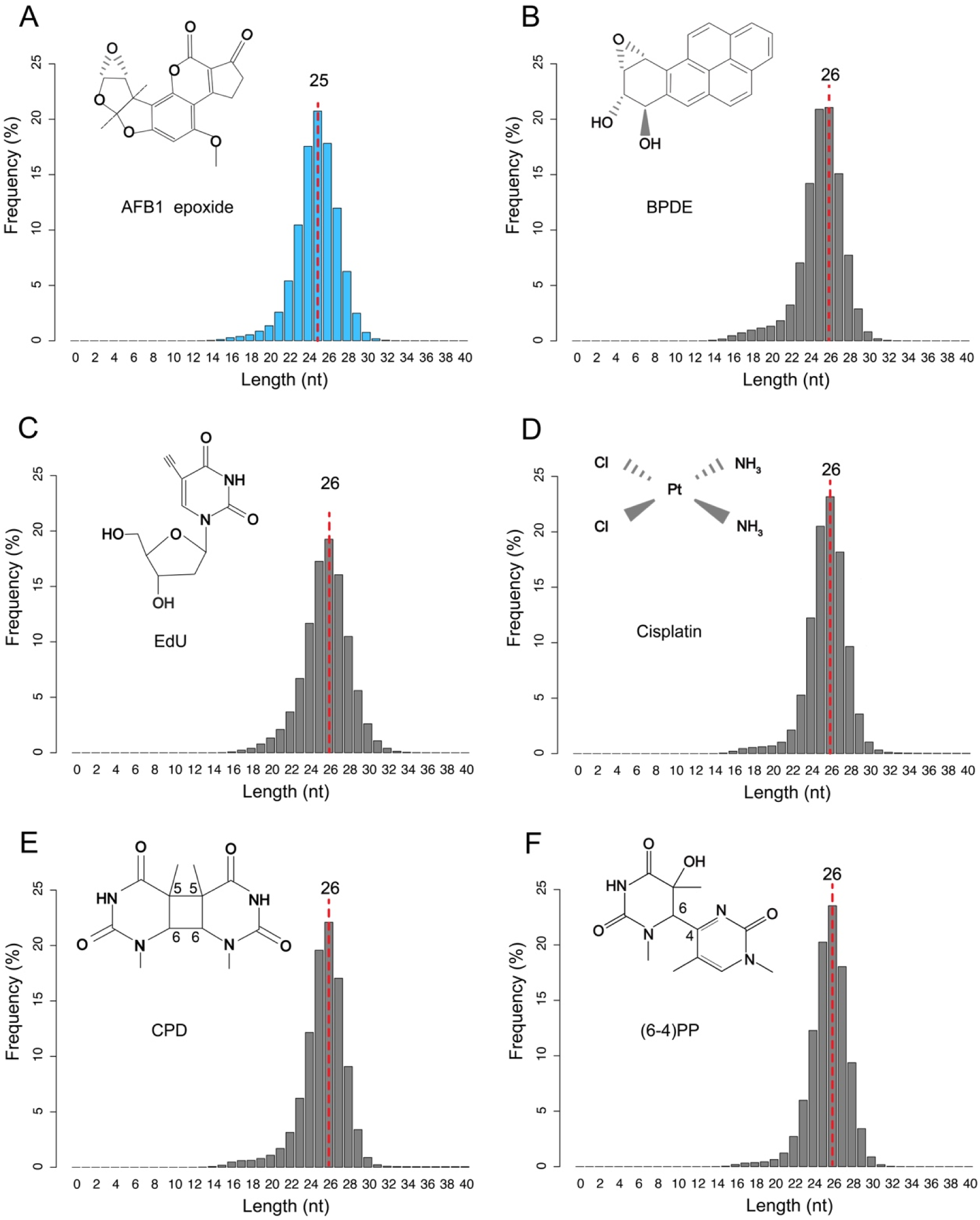
Length distributions of excision products obtained through tXR-seq for AFB1-dG (A) and BPDE (B), and XR-seq for EdU (C), cisplatin (D), CPD (E), and (6–4)PP (F). Excision products with the highest frequency are indicated by a red dashed line.

### Effect of transcription on AFB1-dG repair

Nucleotide excision repair of DNA lesions throughout the whole genome is affected by various factors such as 3D genome organization, regulatory factor binding, DNA replication, transcription, and nucleosome structure. The effect of transcription on nucleotide excision repair has been extensively studied since the discovery of TCR in human cells (*41*). As aforementioned, AFB1-dG adducts are repair-resistant because of their intercalated conformations in the DNA double helix. Thus, it is conceivable that AFB1-dG adducts are mainly removed through TCR. Indeed, analysis of the AFB1-dG repair profiles for TS and NTS revealed that TS is preferentially repaired over NTS (Fig. 3A). The trend of AFB1-dG repair around the transcription start sites (TSS) and transcription end sites (TES) is in general agreement with previously identified CPD repair in human NHF1 and GM12878 cells (*21, 33*). As expected, AFB1-dG repair for both TS and NTS peaks around TSS and gradually declines towards TES. To visualize the interplay between AFB1-dG repair and transcription throughout the whole genome, we used datasets from AFB1-dG tXR-seq and RNA-seq to generate an AFB1-dG repair map with a resolution extending to the single nucleotide level. Fig. 3B shows the transcription and AFB1-dG repair maps of the *TP53* tumor suppressor gene on chromosome 17. It is evident that the AFB1-dG repair level on the TS of *TP53* is higher than the NTS. The preferential repair on TS can also be seen in its adjacent *WRAP53* gene, which is transcribed in the opposite direction.

**Fig. 3.**
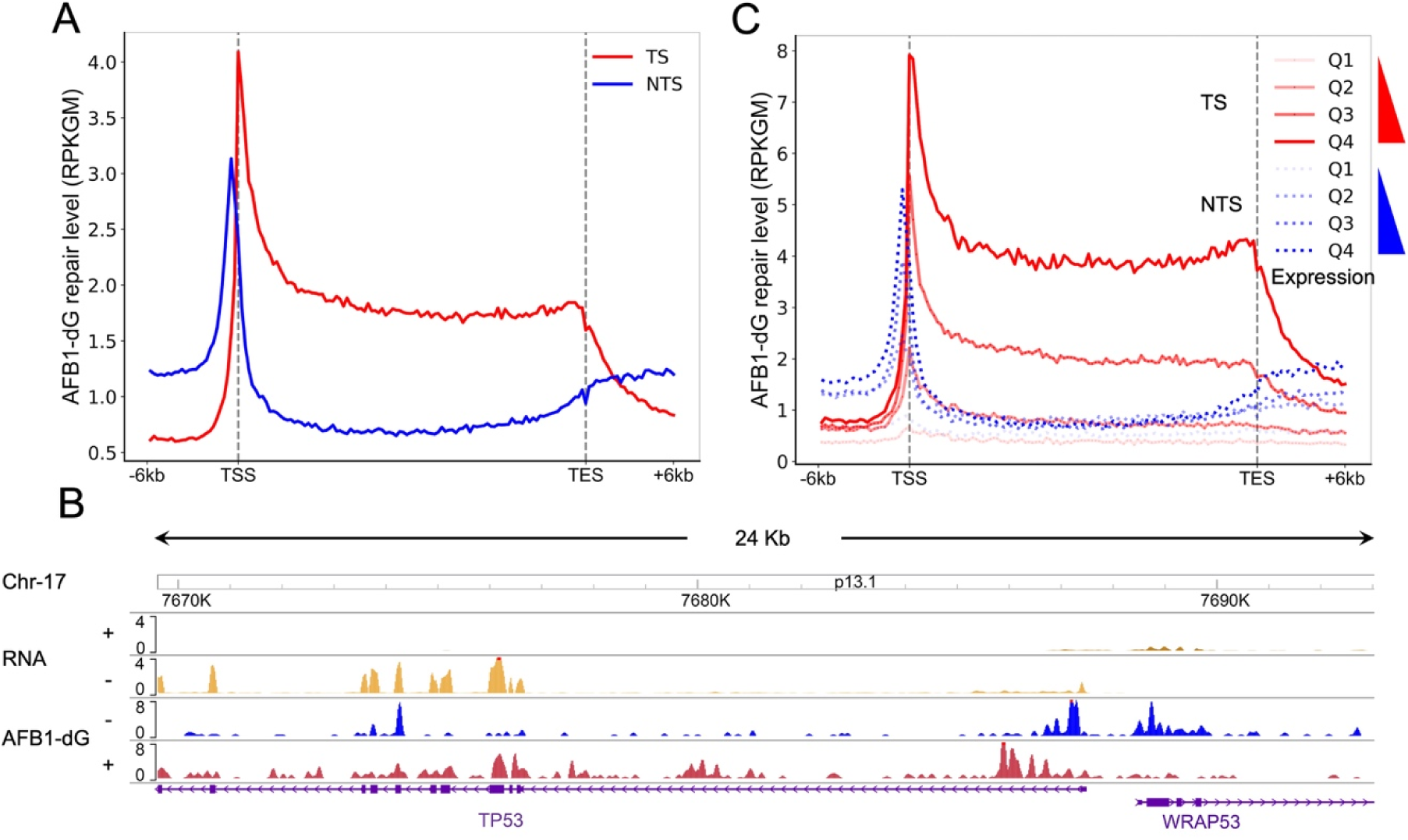
Effect of transcription on AFB1-dG repair. (A) AFB1-dG repair profiles around the transcription start and end sites of 20441 selected genes at 4 h in HepG2 cells. TS and NTS are shown in red and blue, respectively. TS, transcribed strand; NTS, nontranscribed strand. (B) AFB1-dG repair profiles of *TP53* and *WRAP53* genes. “+” in RNA denotes genes transcribed from left to right, while “-” indicates right-to-left transcription. In tXR-seq, “+” signifies plus-strand DNA (5’ to 3’ direction), and “-” denotes minus-strand DNA (3’ to 5’ direction). (C) AFB1-dG repair profiles over 20441 selected genes separated into quartiles based on expression score. Q1 is the lowest expression quartile and Q4 is the highest one.

We next proceeded to investigate the impact of gene expression on AFB1-dG repair. To achieve this, genes were categorized into four quartiles based on their expression levels, ranging from Q1 (the lowest expressed 25% of genes) to Q4 (the highest expressed 25% of genes), and then the average strand-specific repair profiles were plotted for each of these quartiles. A positive correlation for both TS and NTS was observed between AFB1-dG repair and gene-expression levels (Fig. 3C), suggesting that higher gene expression levels are associated with more efficient repair of AFB1-dG bulky adducts.

### Impact of 3D genome organization on AFB1-dG repair

In each human cell, the 23 pairs of chromosomes, approximately 2 meters long if stretched end to end, are packaged into a 5-10 μm diameter nucleus through a hierarchy of compaction. The spatial structures of the human genome are organized at four levels: chromosome territories, A/B compartments, TADs, and chromatin loops (Fig. 4A). Unlike in vitro repair assays conducted with linear DNA, the cellular DNA repair machinery faces the challenge of surmounting the structural barriers created by DNA packaging, which is necessary to gain access to DNA damage for the repair. While various factors, including transcription, DNA replication, histone modifications, circadian clock, and regulatory protein binding, have been found to influence DNA damage and repair (*33, 42–45*), the impact of 3D genome organization on nucleotide excision repair remains largely unknown.

**Fig. 4.**
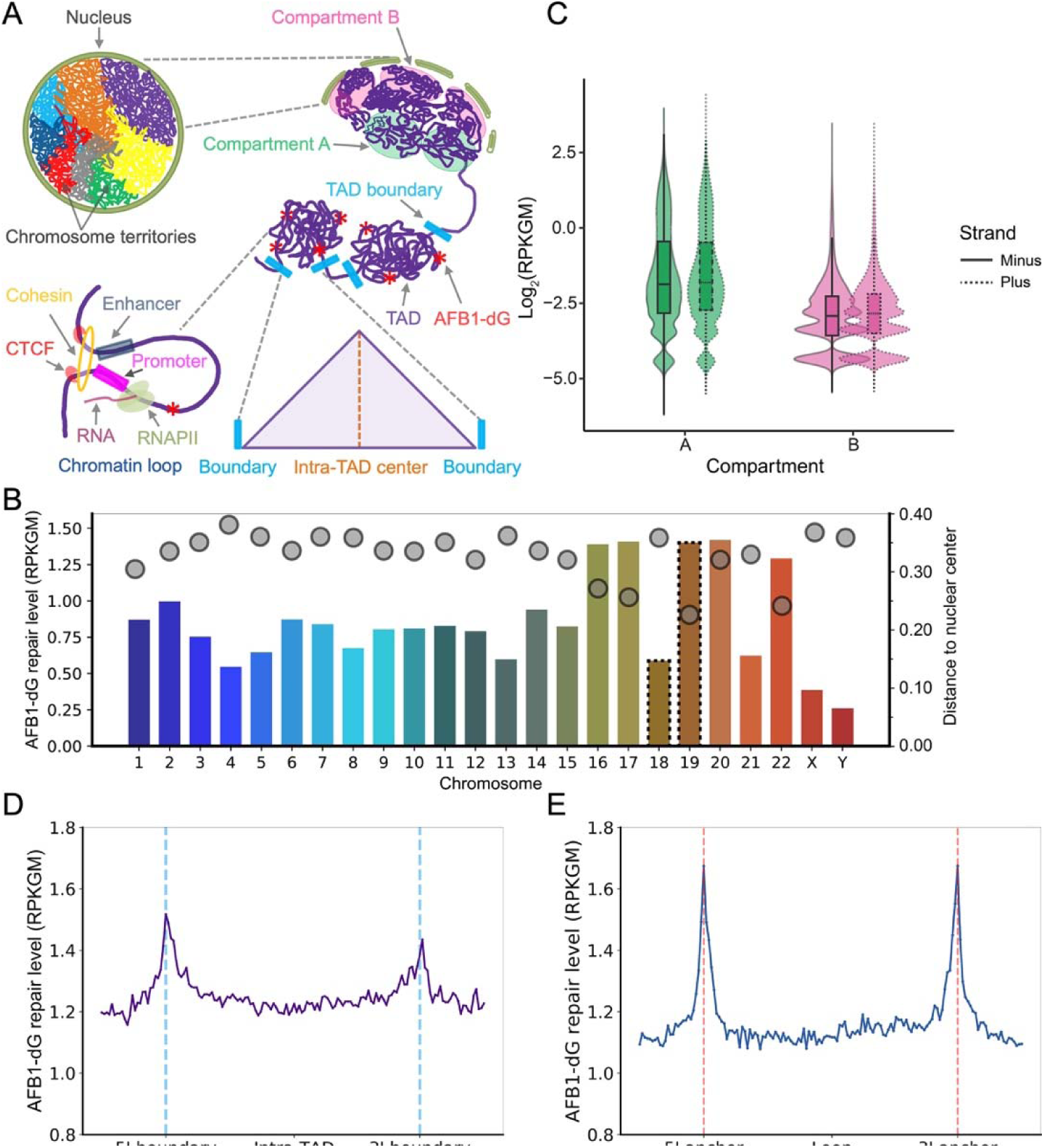
Impact of 3D genome organization on AFB1-dG repair. (A) The hierarchical 3D genome is structured at four levels: chromosome territories, A/B compartments, TADs, and chromatin loops. AFB1-dG DNA adducts within TADs are marked by red stars. (B) Chromosome wise distribution of AFB1-dG repair and chromosome position within the human nucleus. The AFB1-dG repair level on each chromosome is depicted by bar plot. HAS-18 and -19 are highlighted by a dashed line. Gray dots indicate the distance to the nuclear center for each chromosome. (C) Violin plots illustrate AFB1-dG repair levels in A (green) and B (pink) compartments. Plus and minus strands are represented by solid and dashed lines, respectively. The solid band within the box represents the median. (D) Average AFB1-dG repair level over 4311 nonoverlapping TADs. 5’ and 3’ boundaries are denoted by blue dashed lines. (E) Average AFB1-dG repair level across 2989 nonoverlapping chromatin loops that are less than 200 kb. 5’ and 3’ anchors are denoted by red dashed lines.

To explore the impact of chromosome territory positioning on nucleotide excision repair, we calculated the AFB1-dG repair levels on the 23 chromosomes and compared them with the DNA FISH data (*46*). As shown in Fig. 4B, our results indicate that AFB1-dG repair exhibits a non-uniform distribution across all the chromosomes. Notably, repair levels tend to be higher for chromosomes positioned closer to the nuclear center. This trend is evident in chromosomes 18 and 19 (HSA-18 and -19), which are situated in the nuclear periphery and center, respectively. The repair level of gene-rich HSA-19 is approximately 3-fold higher than that of gene-poor HAS-18.

The A compartments are localized within the interior of the nucleus and represent regions of open and expression-active chromatin marked by histone modifications such as H3K4me3 and H3K27ac. In contrast, the B compartments lie at the nuclear periphery and are associated with closed and expression-inactive chromatin, marked by repressive histone markers like H3K9me3 (Fig. 4A). To investigate how genome compartmentalization affects AFB1-dG repair, we analyzed the repair levels in A/B compartments identified from in situ Hi-C experiments in HepG2 cells (*47*). As aforementioned, AFB1-dG adducts are repaired mainly through TCR subpathway, it is plausible to predict that AFB1-dG adducts in the A compartments are repaired faster than those in B compartments. Indeed, the average AFB1-dG repair levels on the two DNA strands in A compartments surpass those observed in B compartments (Fig. 4C).

To determine the impact of TADs on AFB1-dG repair, we generated the average AFB1- dG tXR-seq signal relative to TADs defined with Hi-C data from HepG2 cells. Remarkably, our findings reveal a notable enrichment of average AFB1-dG repair levels around both the 5’ and 3’ boundaries of non-overlapping TADs (Fig. 4D and fig. S4A). It’s noteworthy that the repair level at the 5’ boundaries is higher than the 3’ boundary, and the repair levels in the intra-TAD center are comparable to inter-TAD regions. Given that both TADs and chromatin loops are established through the loop extrusion process and dependent on the binding of CTCF and cohesion, it is reasonable to anticipate that chromatin loops have a similar repair pattern to that observed within TADs. As expected, our analysis demonstrates a significant enrichment of average AFB1-dG repair signals around the two anchors of non-overlapping chromatin loops, all of which are less than 200 kb in size (Fig. 4E). In contrast to TADs, the repair signals at 5’ and 3’ loop anchors exhibit similar levels.

The impacts of genome compartmentalization, TADs, chromatin loops, and transcription on AFB1-dG repair are further illustrated in Fig. 5. In Fig. 5A, the Hi-C heatmap of chromosome 17 at a resolution of 25 kb is displayed using the 3D Genome Browser. Additionally, data from AFB1-dG tXR-seq, Hi-C, RNA-seq, and ChIP-seq for CTCF, the cohesion subunit RAD21, and three histone marks (H3K4me3, H3K27ac, and H3K9me3) are visualized using the WashU Epigenome Browser. As apparent from the data, AFB1-dG repair signals are notably higher within the A compartment (shaded in green) compared to the B compartment (shaded in pink). Meanwhile, ChIP-seq signals for active histone marks (H3K4me3 and H3K27ac) within the A compartment significantly surpass those in the B compartment. Zooming into the yellow-shaded region in Fig. 5A, we identified a TAD spanning 200 kb, containing the NXN gene, where AFB1-dG repair signals are enriched around the 5’ and 3’ boundaries (shaded in light blue). Within this TAD, a loop, denoted by a blue circle, is evident, featuring two anchor regions highlighted in red (Fig. 5B). Notably, AFB1-dG repair signals are higher around these anchor regions, mirroring the heightened ChIP-seq signals for CTCF and RAD21. Furthermore, we observed that AFB1-dG repair on the TS of the NXN gene substantially exceeds that on the NTS, indicating the substantial impact of transcription on the repair process.

**Fig. 5.**
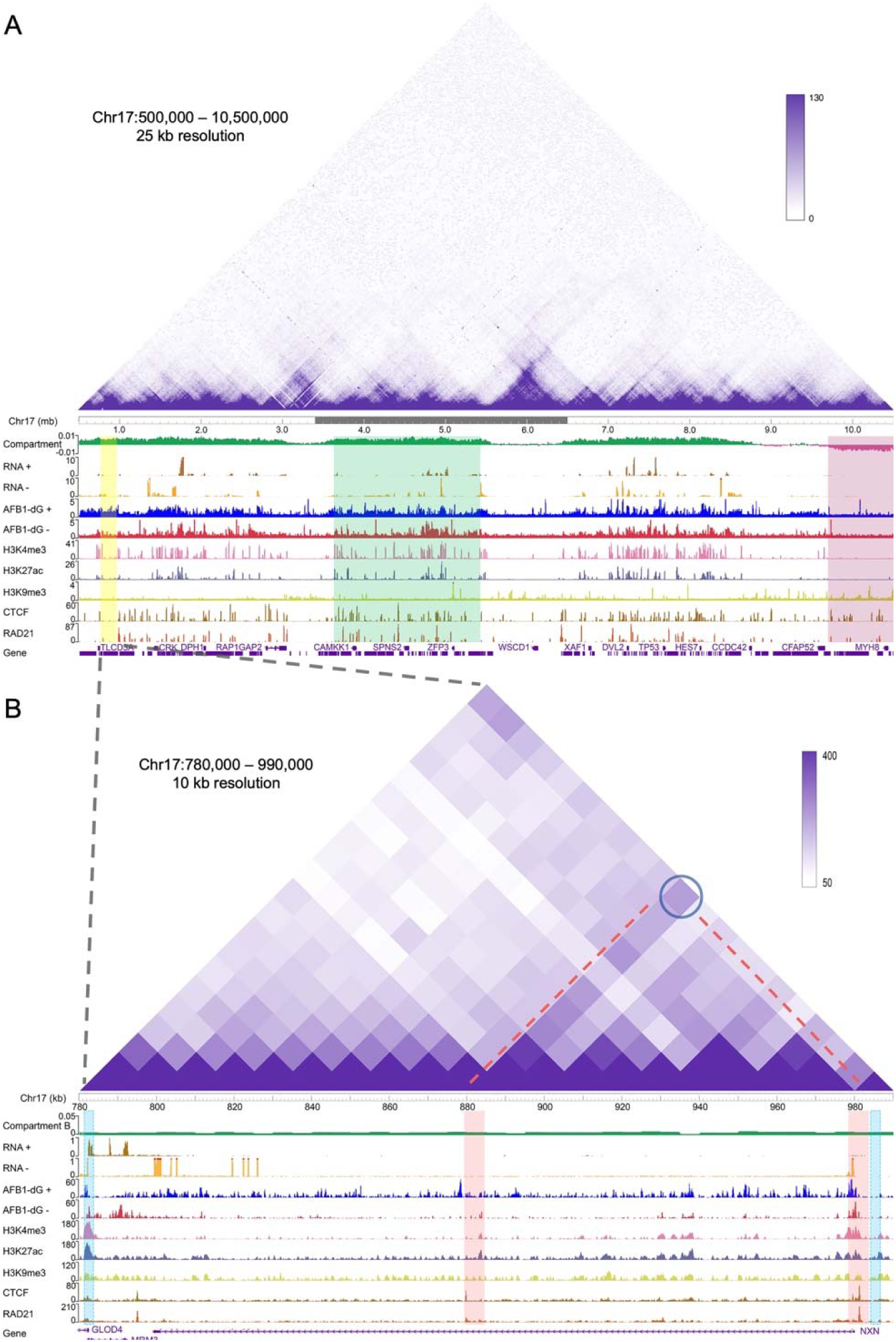
Screenshots of AFB1-dG repair for chromosome 17 across A/B compartments, TADs, and chromatin loops. (A) Hi-C heatmap at 25-kb resolution covering a 10 mb region on chromosome 17. A and B compartments are shown in green and pink, respectively. AFB1-dG tXR-seq data and public RNA-seq data, ChIP-seq data for histone marks (H3K4me3, H3K27ac, and H3K9me3), CTCF, and RAD21 are shown below the heatmap. The selected region is highlighted in yellow. (B) Zoomed view of AFB1-dG repair over the highlighted 210 kb region in Fig. 5A. TAD and anchor boundaries are highlighted in light blue and red, respectively. The representative chromatin loop is indicated by a dark blue circle and red dashed line.

Altogether, our findings suggest that the hierarchical levels of 3D genome organization have a discernible impact on nucleotide excision repair of AFB1-dG adducts, depending on their structural attributes. Evidently, AFB1-dG adducts residing on chromosomes proximal to the nuclear center undergo expedited repair processes. The A compartments, associated with an open chromatin landscape, are preferentially targeted for repair, in contrast to the relatively less accessible B compartments. Furthermore, our results indicate an accelerated repair process occurring at both the boundaries of TADs and the anchor regions of chromatin loops. These findings underscore the impact of 3D genome organization on the nucleotide excision repair machinery in maintaining genomic integrity.

### Effects of size and strength of TADs and chromatin loops on AFB1-dG repair

TADs, characterized by an elevated contact frequency within them, are conserved not only across diverse cell types but also between different species (*48*). These structural units vary in size, ranging from 40 kb to 3 Mb, with a median length of 185 kb (*49*). Many TADs show “corner peaks” in their Hi-C heatmaps, indicating the presence of chromatin loops. These “corner peaks” mark the anchor points of these chromatin loops. To investigate the impact of TAD size on AFB1-dG repair, we conducted an analysis of the AFB1-dG repair levels within non-overlapping TADs as a function of TAD size. As illustrated in Fig. 6A, our findings reveal a notable decrease in repair with increasing TAD size. To examine the effect of TAD strength on AFB1-dG repair, we categorized TADs into quartiles based on their strength and plotted the average repair levels for each quartile (Fig. 6B). We observed a positive correlation between TAD strength and AFB1-dG repair levels. Notably, within the highest strength quartile (Q4), the repair level in the intra-TAD region is much higher than that in the inter-TAD region. Conversely, within the lowest strength quartile (Q1), the reverse pattern is observed, suggesting that strong TAD strength may promote the repair process. We then assessed the impact of chromatin loop size and strength on AFB1-dG repair, employing a similar analytical approach. We found that loop size negatively correlates with repair, while loop strength is positively associated with repair, which mirrors the trends observed within TADs (Fig. 6, C and D, and fig. S4B). Subsequently, we extended our analysis to evaluate the impact of loop size and strength identified from CTCF ChIA-PET data on AFB1-dG repair. Our findings revealed a similar trend, with loop size negatively correlating with repair (fig. S4C). However, we observed that the strength of loops identified from the CTCF ChIA-PET experiment did not exhibit a noticeable effect on AFB1-dG repair (fig. S4D).

**Fig. 6.**
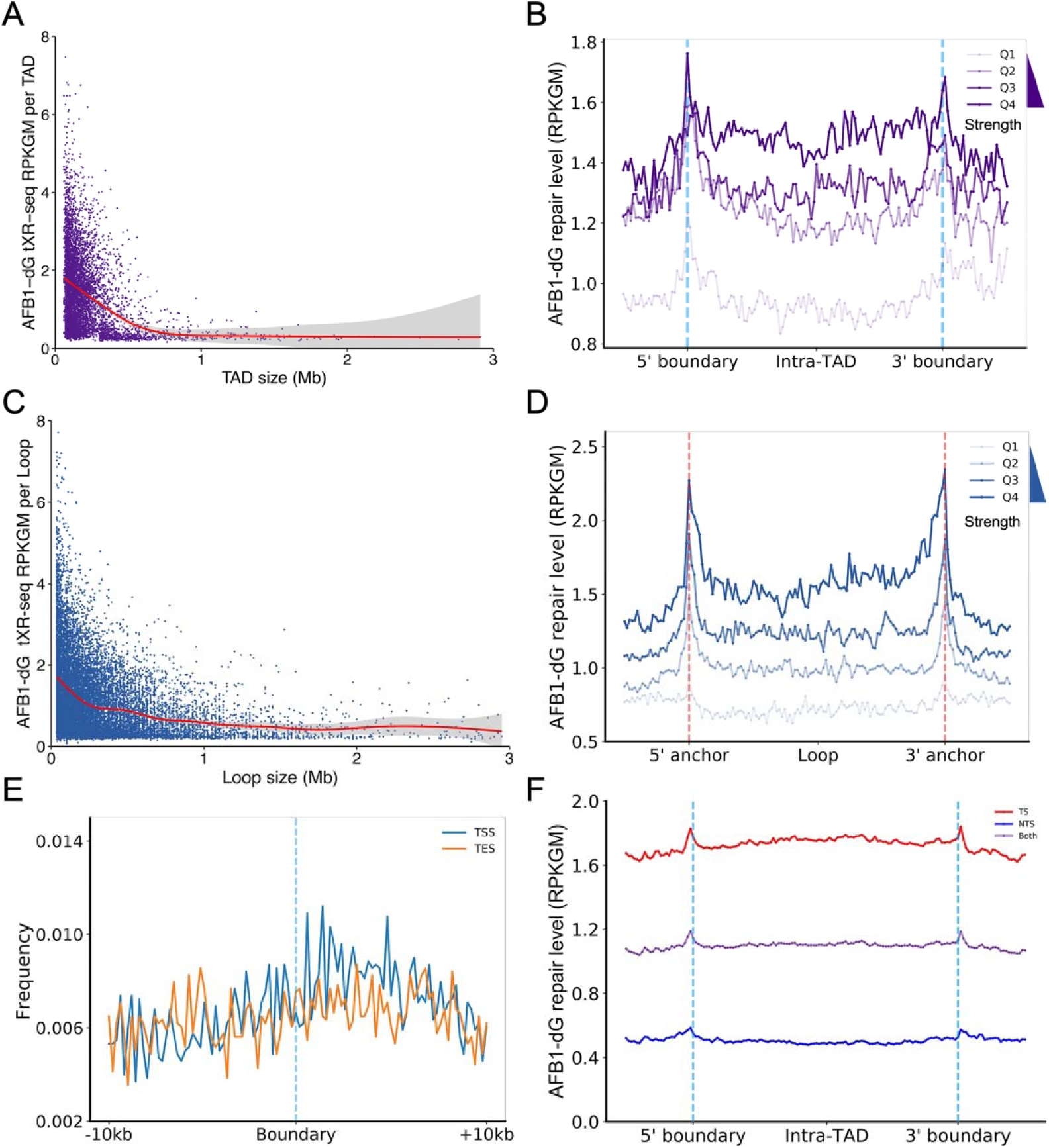
Effects of size and strength of TADs and chromatin loops on AFB1-dG repair. (A) A scatter plot illustrates AFB1-dG repair levels per TAD relative to TAD size. There are a total of 6183 TADs included in this analysis. (B) AFB1-dG repair profiles are displayed across 4311 nonoverlapping TADs, categorized into quartiles based on TAD strength scores, ranging from Q1 (lowest strength) to Q4 (highest strength). (C) A scatter plot shows AFB1-dG repair levels per chromatin loop relative to loop size. The total number of chromatin loops shown here is 14445. (D) AFB1-dG repair profiles are shown across 2989 nonoverlapping chromatin loops (length < 200 kb), separated into quartiles based on loop strength scores, ranging from Q1 (lowest strength) to Q4 (highest strength). (E) Frequency distributions of TSS and TES surrounding boundary regions of 4311 nonoverlapping TADs. (F) AFB1-dG repair profiles for TS and NTS over 4311 nonoverlapping TADs.

Our results revealed elevated levels of AFB1-dG repair within regions encompassing TAD boundaries and loop anchors. We hypothesized that this phenomenon may be attributed to the elevated levels of transcription activity within those regions. Indeed, we examined the distribution of TSS and TES around TAD boundaries and found an enrichment of TSS frequencies within boundary regions (Fig. 6E). Notably, TSS signals displayed a decrease in the center of both 5’ and 3’ boundaries. Around the 5’ boundary region, TSS frequency was notably high, while TSS signals exhibited an increase only downstream of the 3’ boundary (fig. S4, D and E). Further analysis of the average repair profiles for the TS and NTS across non-overlapping TADs revealed that TS repair levels were approximately four-fold higher than NTS repair levels (Fig. 6F). Interestingly, within both the 5’ and 3’ boundary regions, NTS repair levels were also elevated. This suggests that factors beyond transcription-associated factors, such as histone modifications and chromatin-binding proteins, may contribute to the elevated repair observed on the NTS within boundary regions.

## DISCUSSION

In the liver cell, AFB1, the most common and potent carcinogen, is metabolized to AFB1-8,9-epoxide which intercalates in DNA to form AFB1-dG bulky adducts (AFB1-N^7^-dG and AFB1-FAPY-dG). In addition, the reactive oxygen species produced during AFB1 metabolism induce lipid peroxidation and its byproducts, such as acetaldehyde and crotonaldehyde, can form bulky DNA adducts as well (*13*). The NER pathway removes AFB1- dG adducts through the dual-incision mechanism to maintain genome integrity. If left unrepaired, AFB1-dG bulky adducts can cause mainly G to T transversions such as the mutation hotspot at codon 249 of the *TP53* gene found in HCC patients. The whole human genome is hierarchically packaged into chromatin on several levels inside the nucleus. The 3D genome organization plays a critical role in many cellular processes including DNA replication, transcription, DNA damage, and repair. To understand the molecular mechanism of AFB1 hepatocarcinogenesis, it is important to delineate the molecular details of AFB1-induced DNA damage formation and subsequent repair events across the entire human genome. Previous studies focused on specific genes such as *TP53* gene by using conventional radioactive labeling-based methods including LMPCR (*7, 50*). However, those studies that interrogated AFB1-induced DNA damage and repair in linear DNA sequences lack the information of 3D genome organization. It remains unknown how AFB1-dG adducts are repaired at genome-wide level and how 3D genome organization affects the AFB1-dG repair. Here, for the first time, we mapped the AFB1-dG repair across the whole genome with single nucleotide resolution and revealed a heterogeneous AFB1-dG repair landscape across the four different genome organization levels: chromosome territories, A/B compartments, TADs, and chromatin loops.

Our adaptation of tXR-seq method has enabled us to create a genome-wide repair map of AFB1-dG adducts with single nucleotide resolution in humans. As expected, we observed the preferential repair on TS of *TP53* gene, suggesting the repair resistant AFB1-dG adducts, like CPD lesions, are primarily eliminated through TCR subpathway. In our analysis, we revealed a distinct length distribution pattern of excision products released during AFB1-dG repair. The predominant excision product length is only 25 nt long, which is 1 nt shorter than previously identified excision products resulting from the repair of a variety of DNA bulky adducts caused by substances such as BPDE, EdU, cisplatin, and UV. We hypothesize that this unique pattern might be attributed to the relatively large size of AFB1-dG adduct itself. This size discrepancy could potentially pose challenges for the TFIIH complex to cover the same length of DNA fragment containing AFB1-dG compared to other smaller bulky adducts, such as CPD and (6–4)PP. In this context, there might be two scenarios: 1) the XPF and XPG may execute the dual incision flanking the AFB1-dG adduct in a manner identical to their approach with other bulky adducts. The consistent dual incision pattern generates excision products of the same size, resulting in one extra nucleotide dangling out of the TFIIH complex. Exonucleases such as TREX1 will then quickly degrade the single extra nucleotide; 2) the XPF and XPG may employ the dual incision strategy one nucleotide shorter when dealing with the AFB1-dG adduct compared to their reactions to other smaller bulky adducts. This scenario gains plausibility from the analysis of nucleotide frequency within 24-mers, 25-mers, 26-mers and 27-mers released during AFB1-dG repair, which suggests that dual incision occurs approximately 18-20 nt 5’ and 5-6 nt 3’ to the lesion. In contrast, for other smaller DNA lesions, the dual-incision sites are typically at 19-21 nt 5′ and 5-6 nt 3′ to the lesion (*51, 52*). However, further work is required to validate the precise dual incision reactions for repairing AFB1-dG adducts in humans.

Our in-depth analysis of AFB1-dG repair within the context of 3D genome organization provided several critical insights into the complex mechanisms underlying the dynamics of AFB1-dG repair. Integrating multiple datasets, including tXR-seq, DNA FISH, Hi-C, and ChIA-PET, we revealed that AFB1-dG repair is heterogeneous across the chromosome territories, A/B compartments, TADs, and chromatin loops. We observed the dependency of AFB1-dG repair on the spatial positioning of chromosomes within the nuclear architecture. Chromosomes located closer to the nuclear center, such as HSA-19, exhibited faster repair compared to those situated in the nuclear periphery, like HAS-18. This finding implies a spatial regulation of repair activity within the cell nucleus. Furthermore, we identified the effect of A/B compartments on AFB1-dG repair. The A compartments, characterized by active and euchromatin regions, displayed higher AFB1-dG repair levels relative to the B compartments, which are associated with inactive and heterochromatin regions. In our previous work, we demonstrated that CPD and (6–4)PP repair super-hotspots are enriched in frequently interacting regions (FIREs) and super-enhancers (*53*). Here, we identified the enrichment of AFB1-dG repair in the vicinity of TAD boundaries and chromatin loop anchors. Interestingly, we found the strength of TADs and loops are positively associated with repair levels, while there was a negative correlation with the size of these genomic structures. These results suggest that genomic regions with robust transcriptional activity are favored for repair through the TCR subpathway. Meanwhile, epigenetic regulation, particularly histone modifications that maintain a euchromatin state, might contribute to the high global repair on NTS within regions exhibiting high TCR.

Although TADs are highly conserved fundamental units of 3D genome organization, their structures are dynamic in cycling mammalians cells (*54*). When exposed to DNA-damaging agents, the DNA damage response and subsequent repair processes can potentially impact the 3D genome organization. Notably, it has been documented that TAD boundaries are strengthened, which is attributed to increased recruitment of the architectural protein CTCF after ionizing radiation (*55*). Previous studies have demonstrated that CTCF recruitment to double-strand break sites promotes homologous recombination repair and cohesion-mediated loop extrusion on either side of the break contributes to the formation of γH2AX and DNA damage response foci (*56–58*). It is plausible that AFB1 exposure may induce similar TAD boundary strengthening to facilitate efficient nucleotide excision repair.

Recent studies have shown that chromosomes located near the nuclear periphery are more susceptible to damage from UV radiation (28, 57, 58). These studies have indicated that both CPD and (6–4)PP are enriched in heterochromatin regions, suggesting that peripheral genome organization serves as a protective shield for nuclear center regions against UV-induced damage. Beyond the alteration of 3D genome organization in response to DNA damage, the distribution of DNA damage within various layers of 3D genome organization may also affect the repair process.

To comprehensively dissect the interplay between 3D genome organization and the repair of AFB1-dG adducts, further investigations into the alterations of 3D genome organization in response to AFB1 exposure are needed. Additionally, profiling the distribution of AFB1-dG adducts within the 3D genome structure is required. These future studies will enhance our understanding of the dynamic relationship between DNA damage, repair, and 3D genome architecture, offering valuable insights into the mechanisms underlying AFB1-induced HCC.

## MATERIALS AND METHODS

### Cell line and culture conditions

The human HepG2 cell line was purchased from the America Type Culture Collection (ATCC). AFB1 (Cat. No. A6636) was obtained from MilliporeSigma. HepG2 cells were cultured in Dulbecco’s Modified Eagle Medium (DMEM) supplemented with 10% FBS at 37 °C in a 5% CO_2_ humidified chamber.

### UVC irradiation and AFB1 treatment

HepG2 cells were cultured until they reached approximately 80% confluence, after which they were subjected to UVC irradiation (20 J/m^2^) using a GE germicidal lamp emitting primarily 254-nm UV light. Following irradiation, the cells were cultured at 37°C for 0.5 hours before conducting an *in vivo* excision assay. For aflatoxin treatment, an AFB1 stock solution (4 mM) was added to the HepG2 cell culture medium in a cell culture flask, resulting in a final concentration of 40 µM. The cells were cultured for 4 hours, with gentle shaking every 30 minutes to ensure cell suspension in the medium.

### Antibodies, TLS DNA polymerase, and oligonucleotides

The following antibodies were used in this study: anti-XPB (Santa Cruz, sc293), anti-XPG antibody (Santa Cruz, sc13563), rabbit anti-mouse IgG (Abcam, ab46540), anti-aflatoxin monoclonal antibody (6A10) (Thermo Fisher, MA1-16885), and anti-AFB1 (AFA-1) (Abcam, ab1017). Sulfolobus DNA polymerase IV (Cat. No. M0327) and DNA polymerase ζ (Cat. No. 51) were purchased from NEB and Enzymax, respectively. Oligonucleotides used for adaptor ligation and PCR amplification of the sequencing library were the same as described previously (*21*).

### *In vivo* excision assay

The *in vivo* excision assay procedure is the same as previously described (*59*). Briefly, after AFB1 treatment, cells were lysed by using the Hirt lysis method. The supernatant was subject to IP with anti-TFIIH and anti-XPG antibodies for capturing the excision products released during nucleotide excision repair. The isolated excision products were then 3’ end-labeled with ^32^P- Cordycepin and resolved on a 10% sequencing gel.

### AFB1-dG tXR-seq library preparation and sequencing

AFB1-dG tXR-seq libraries were prepared as described in our previous study (*33*). Briefly, the excision products were captured by TFIIH/XPG IP and treated with carbonate-bicarbonate buffer (pH 9.6) to convert the AFB1-N^7^-dG to the ring-opened AFB1-FAPY-dG adducts. Following adaptor ligation, the adaptor-ligated excision products were further purified by IP with anti-aflatoxin monoclonal antibody. Then sulfolobus DNA polymerase IV was used to bypass the AFB1-FAPY-dG adduct during the primer extension step of tXR-seq. After purification of the primer extension products, Kapa Hotstart Readymix was used for amplifying the library. All tXR-seq libraries were sequenced on the Illumina HiSeq 2500 platform.

### Data processing and visualization

Two biological replicates of AFB1 tXR-seq were used for the analysis. The adaptors were trimmed by using BBduk and reads longer than 50 mers were filtered out for analysis. PCR duplicates were removed by the FASTX-Toolkit. Refined reads were aligned to the human reference genome GRCh38 by using the bowtie tool (*60*) with arguments: -x -q --nomaqround -m 4 -v 3 --tryhard --strata --best -p 4 --seed=123. The aligned reads were normalized with the sequencing depth and visualized with Integrative Genomics Viewer (*61*).

### Data collection

XR-seq datasets used for length distribution analysis were downloaded from the Gene Expression Omnibus with accession numbers: GSE97675 (BPDE), GSM6222398 (EdU), GSE82213 (cisplatin), GSE67941 [CPD and (6–4)PP]. The processed *in-situ* Hi-C datasets from the HepG2 cell line were downloaded from the ENCODE consortium (ID: ENCSR194SRI). The processed CTCF-mediated ChIA-PET data from HepG2 cells was downloaded from ENCODE (ID: ENCSR411IVB).

### Length distribution and nucleotide frequency analysis

The length distributions for all reads and single nucleotide frequencies for selected excision products from tXR-seq and XR-seq were calculated using in-house custom scripts.

### Average AFB1-dG repair profile analysis

The AFB1-dG repair level for each selected region such as chromosome, A/B compartment, TAD, loop, gene, TS, and NTS was counted and normalized to RPKM (Reads Per Kilobase per Million mapped reads). As AFB1-8,9-epoxide preferentially binds to the N^7^ position of guanine, the GC content within each selected region may cause bias in our analysis. We then used the total number of Gs in the selected region for the normalization, which we termed “RPKGM” shown in the following equation:

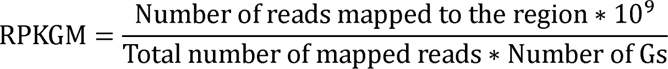

For analysis of the effect of transcription on AFB1-dG repair, the gene list file of the GRChg38 genome assembly was downloaded from the USCS genome browser. Genes with an expression score greater than 300 and a length greater than 3 kb were selected. Among those genes, those with another neighboring transcript within the ±6kb vicinity were removed. The selected 20441 genes were divided into 100 bins and their ±6kb regions were divided into 50 bins. Then, they were intersected with the mapped read counts of both positive and negative strands to get the read counts for both TS and NTS. The numbers of tXR-seq reads in each bin were calculated and normalized to get RPKGM values.

For the chromosome wise analysis of AFB1-dG repair, we calculated the total number of Gs for each chromosome from human reference genome GRCh38 and intersected each chromosome with AFB1-dG tXR-seq mapped reads to get the total number of mapped reads. Then, we obtained the AFB1-dG repair level based on the above RPKGM equation. For the A/B compartments analysis, we downloaded A/B Compartments (ID: ENCFF091UMY) of the HepG2 cells from the ENCODE. We separated each compartment into three categories: A, B, and conserved compartments, based on the Principal Component Analysis value. Bedtools was used to intersect the A/B compartments with AFB1-dG tXR-seq mapped reads of both TS and NTS to calculate the repair level in A/B compartments.

In the analysis of TAD effect on AFB1-dG repair, TADs data (file ID: ENCFF018XKF) from the HepG2 cell line was used for the analysis. Overlapping TADs were removed from further analysis, and the resulting nonoverlapping TADs (4311 in total) were intersected with AFB1-dG tXR-seq reads using bedtools (*62*). Read counts were calculated at each TAD and were normalized by the RPKGM equation. Additionally, the selected TADs were divided into quartiles (*Q1 to Q4*) based on the TAD strength score and intersected with the AFB1-dG tXR-seq reads to calculate the number of read counts in the TAD boundaries and intra-TAD regions for each quartile. For TAD boundary analysis, we defined the boundary of each TAD by extending the start and end to four different window sizes ±1kb, ±5kb, ±10kb, and ±50kb and created the new sub-starts and sub-ends of boundary regions. The intra-TAD regions were defined as follows: we initially calculated the center (mid-point) of each TAD, and then extended these center points on both sides using the same methodology as that employed for extending the window sizes at TAD boundaries.

### Statistical Analysis and Visualization

Statistical calculations such as the Wilcoxon test was performed using R-Packages. The ggplot2 package was used for visualization and plotting in R-Studio. Hi-C heatmaps were visualized by WashU Epigenome Browser (http://epigenomegateway.wustl.edu/browser/).

## ACKNOWLEDGMENTS

We thank Dr. Christopher Selby and Dr. Laura Lindsey-Boltz for their proofreading assistance. We also thank all the members of the Li laboratory for helpful discussion and proofreading.

## Author contributions

Conceptualization: A.S. and W.L. Methodology: Y. W., M.M.A and W.L. Investigation: Y. W., M.M.A and W.L. Visualization: Y. W., M.M.A and W.L. Supervision: A.S. and W.L. Writing (original draft): W.L. Writing (review and editing): Y. W., M.M.A, A.S. and W.L.

## Data and materials availability

The raw AFB1 tXR-seq data in this study were deposited in GEO (accession no. GSE243425). Scripts used in this article are available at https://github.com/wentao-li/AFB1_3D_Genome.

