## Supplementary Materials for "Nucleotide Excision Repair of Aflatoxin-induced DNA Damage within the 3D Human Genome Organization"

Yiran Wu et al.

Supplementary Figures 1-4

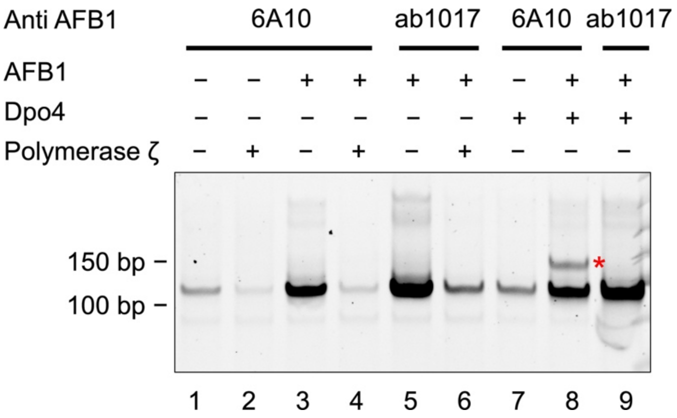


**Fig. S1.** AFB1-dG tXR-seq library preparation involved the use of different anti-AFB1 antibodies (6A10 and ab1017) and translesion synthesis DNA polymerases (Dpo4 and polymerase ζ). The libraries were analyzed through 10% native polyacrylamide gel electrophoresis, where PCR products containing inserts were identified and marked by a red star.

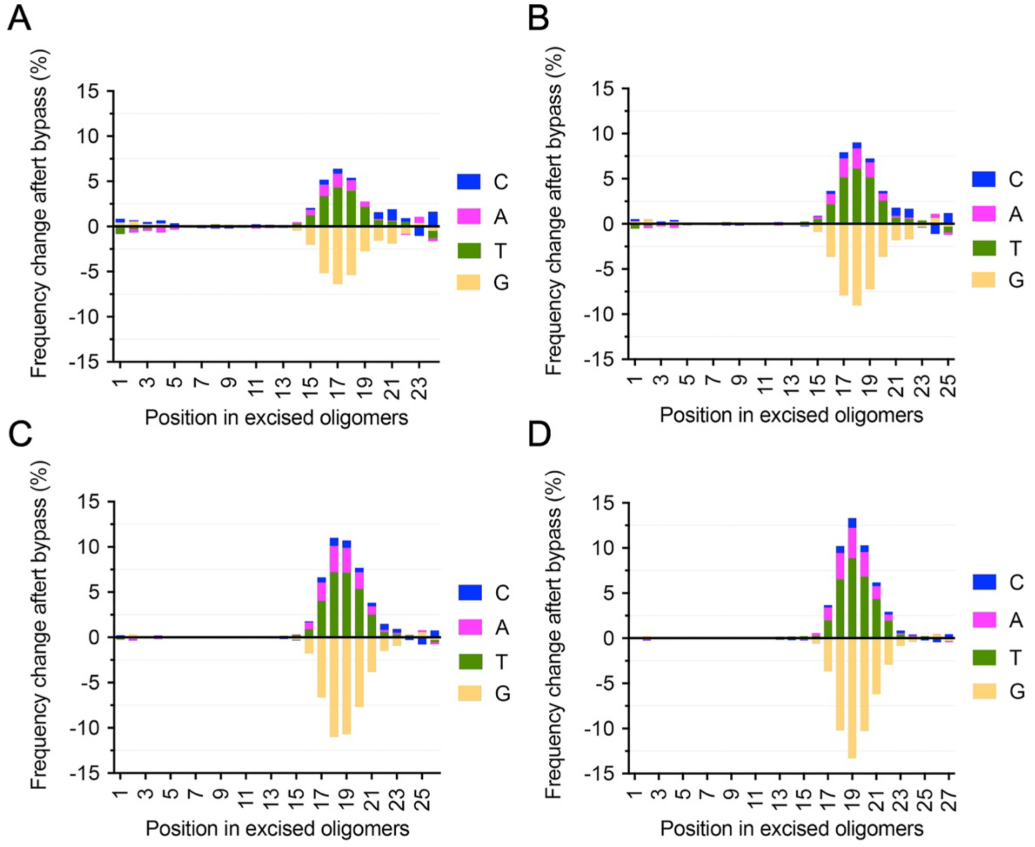


**Fig. S2.** Dpo4 primarily induces G to T transversions. The analysis focuses on frequency changes following bypass in excision products of different lengths, including 24-mers (A), 25-mers (B), 26-mers (C), and 27-mers (D). After bypass read sequences are extracted from the human reference genome after alignment and compared their nucleotide frequencies with those of the raw reads.

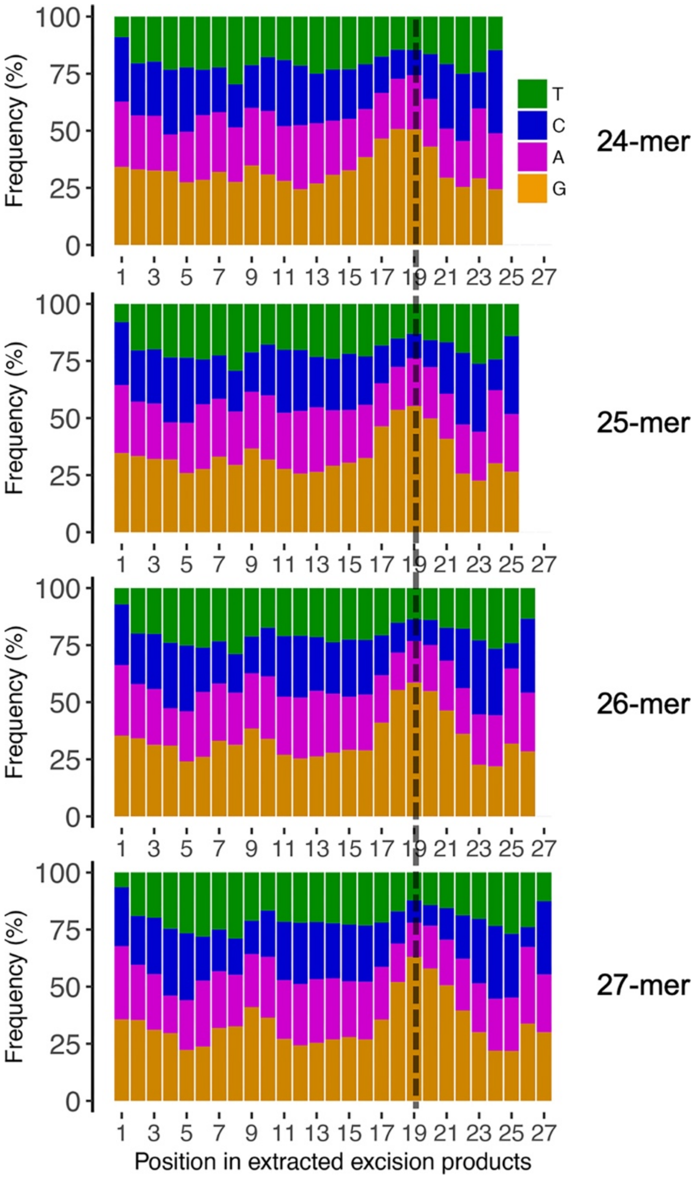


**Fig. S3.** Single-nucleotide frequencies obtained through AFB1-dG tXR-seq extracted reads, covering sequences ranging from 24-mers to 27-mers. Notably, the enrichment of Gs at position 19 is highlighted by a black dashed line.

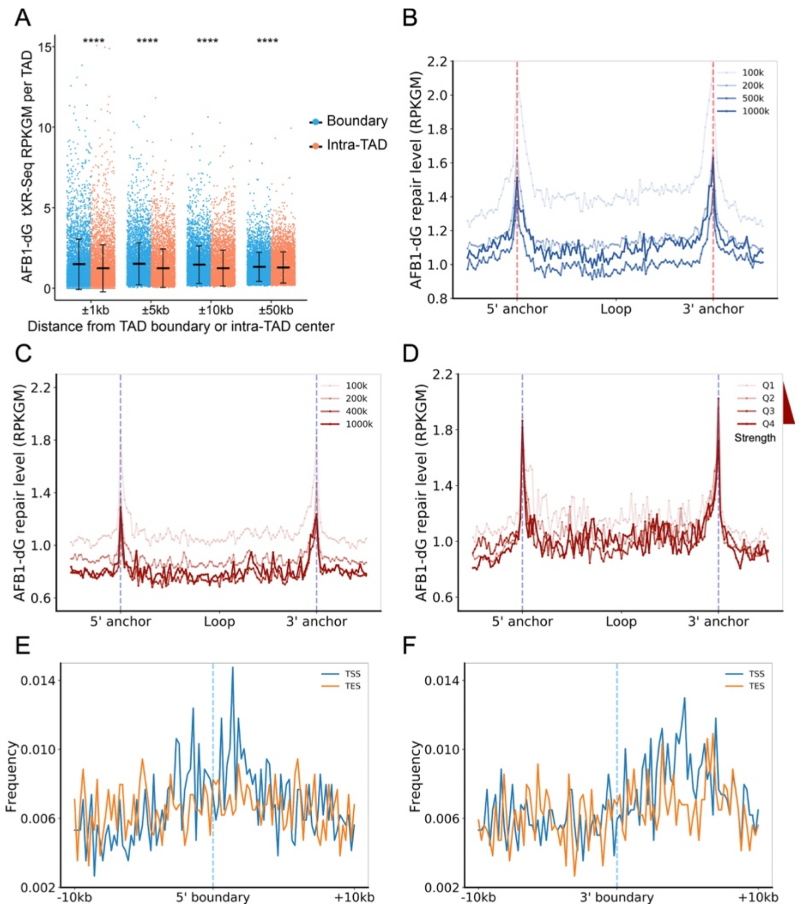


**Fig. S4.** Enrichment of AFB1-dG repair and transcription activity around both TAD boundaries and chromatin loop anchors. (A) AFB1-dG repair levels within extended regions (1kb, 5kb, 10kb, and 50kb) both upstream and downstream of TAD boundaries. Mean values with standard deviation (SD) are indicated with a black error bar. Statistical comparisons between mean values from both groups were conducted, and p-values were calculated using the Wilcoxon rank test. The p-values for ±1kb, ±5kb, and ±10kb are all less than 2e-16, and the p-value for ± 50kb is 5.6e-6. (B) AFB1-dG repair levels across chromatin loops with different lengths from HiC data from HepG2 cells. (C) AFB1-dG repair levels across chromatin loops with different lengths obtained from CTCF ChIA-PET data. (D) AFB1-dG repair levels across 2737 chromatin loops (length < 100 kb) obtained from CTCF ChIA-PET data. The chromatin loops were categorized into quartiles based on loop strength, ranging from Q1 (lowest strength) to Q4 (highest strength). (E) Frequency distributions of TSS and TES surrounding the 5' boundary regions of 4311 nonoverlapping TADs are displayed. (F) Frequency distributions of TSS and TES surrounding the 3' boundary regions of 4311 nonoverlapping TADs are shown.
